# Genomic epidemiology and antifungal-resistant characterization of *Candida auris*, Colombia, 2016-2021

**DOI:** 10.1101/2023.10.20.563341

**Authors:** Elizabeth Misas, Patricia L. Escandon, Lalitha Gade, Diego H. Caceres, Steve Hurst, Ngoc Le, Brian Min, Meghan Lyman, Carolina Duarte, Nancy A. Chow

## Abstract

Since 2016 in Colombia, ongoing transmission of *Candida auris* has been reported in multiple cities. Here, we provide an updated description of *C. auris* genomic epidemiology and the dynamics of antifungal resistance in Colombia. We sequenced 99 isolates from *C. auris* cases with collection dates ranging from June 2016 to January 2021; the resulting sequences coupled with 103 previously generated sequences from *C. auris* cases were described in a phylogenetic analysis.

All *C. auris* cases were of clade IV. Of the 182 isolates with antifungal susceptibility data, 67 (37%) were resistant to fluconazole, and 39 (21%) were resistant to amphotericin B. Isolates predominately clustered by country except for 16 isolates from Bogotá, Colombia, which grouped with isolates from Venezuela. The largest cluster (N=166 isolates) contained two subgroups. The first subgroup contained 26 isolates, mainly from César; of these 85% (N=22) were resistant to fluconazole. The second subgroup consisted of 47 isolates from the north coast; of these, 81% (N=38) were resistant to amphotericin B.

Mutations in the *ERG11* and *TAC1B* genes were identified in fluconazole-resistant isolates, and two amino acid substitutions in PSK76257-(FLO8) and PSK74852 genes were associated with higher minimum inhibitory concentration values for amphotericin B. This work may help identify mechanisms conferring azole and amphotericin B resistance in *C. auris*.

Overall, *C. auris* cases from different geographic locations in Colombia exhibited high genetic relatedness, suggesting continued transmission between cities since 2016. These findings also suggest a lack of or minimal introductions of different clades of *C. auris* into Colombia.

**IMPORTANCE:** *Candida auris* is an emerging fungus that presents a serious global health threat and has caused multiple outbreaks in Colombia. This work discusses the likelihood of introductions and local transmission of *C. auris* and provides an updated description of the molecular mechanisms associated with antifungal resistance in Colombia. Efforts like this tracking genomic variation provide information about the evolving *C. auris* burden that could help guide public health strategies to control *C. auris* spread.

## INTRODUCTION

*Candida auris* is a multidrug-resistant yeast capable of causing outbreaks of invasive infections associated with high mortality rates [1]. Infection and colonization mainly affect patients requiring critical medical care [2, 3]. *C. auris* infections have been reported in over 40 countries [4] since its first description in Japan in 2009 [5].

Analysis of genomic sequences of *C. auris* from global regions showed five major clades. Initially, four clades were identified including clade I (South Asian), clade II (East Asian), clade III (African), and clade IV (South American) [1]. Later in 2019, clade V (Iranian) was reported [6]. Together, these clades show high genetic variation between them and smaller variation within each clade [1]. In Colombia, genomic sequencing revealed all *C. auris* cases were clade IV and suggested ongoing transmission of *C. auris* in multiple cities across the country [7]. Notably, the first clade IV case reported in South America was identified in 2012 in Venezuela, a neighboring country to Columbia [8].

Three classes of antifungal medications are used for *Candida* infections, azoles, polyenes, and echinocandins. Echinocandins are recommended as first-line therapy for invasive candidiasis in many countries [9]. Differences in antifungal resistance have been observed between the clades of *C. auris* [1, 10]. In Colombia, about 35% of *C. auris* isolates have been resistant to fluconazole, and 33% have been resistant to amphotericin B [8]. *C. auris* azole resistance has been reported to increase over time, both in the proportion of isolates resistant to fluconazole as well as an increase in minimal inhibitory concentration (MIC) values [8]. Additionally, a cluster associated with amphotericin B resistance has been identified in the north-coast region of Colombia [7].

Molecular mechanisms associated with or conferring *C. auris* antifungal resistance are increasingly being reported. For azoles, resistance is usually associated with mutations in the *ERG11* gene, a protein-coding gene that encodes lanosterol 14-alpha-demethylase, an enzyme in the ergosterol biosynthesis pathway [11]. These mutations include Y132F, K143R and F444L, which are located close to the active site and the heme cofactor and affect the binding to fluconazole [12, 13]. Additionally, the mutation in *TAC1B* gene has been shown to be associated with fluconazole resistance; *TAC1B* gene encodes a zinc-cluster transcription factor, which activates the ATP-binding cassette (ABC)-type efflux pump-encoding gene CDR1 [14].

For polyenes, molecular mechanisms conferring resistance are largely unknown; previous reports have linked resistance to amphotericin B to overexpression of several mutated *ERG* genes and reduced ergosterol levels [11]. Additionally, Escandon et al., 2019 identified two non-synonymous substitutions associated with high MIC values of amphotericin B. Specifically, substitution S108N in the *FLO8* gene (PSK76257) was observed [7, 15], and a hypothetical protein that encodes a membrane transporter (PSK74852) showed the substitution I139T [7]. The *FLO8* gene is a transcription activating factor that plays a role in hyphal development in *Candida albicans* [16], which may increase its survival in the presence of amphotericin B [17]. The substitution S108N is located in the highly conserved region of LUFS (stands for “LUG/LUH, Flo8, single-strand DNA-binding protein [SSBP]”), which has been identified as a transcriptional regulator proteins of plants, fungi and humans [18]. For echinocandins, a mutation in *FKS1*, the gene that encodes the echinocandin target 1,3-beta-D-glucan synthase, has been widely described as conferring echinocandin resistance [15, 19].

In this study, we investigated genomic epidemiology of *C. auris* cases from 2016 – 2021 in Colombia to obtain an updated description of *C. auris* transmission dynamics and molecular characterization of antifungal resistance.

## MATERIALS & METHODS

### Sample collection

In 2006, the INS published the National Alert on the Emergence of Infections Caused by *Candida auris* [20], which requested that public health laboratories in Colombia send all suspected or confirmed *C. auris* isolates to the Mycology reference laboratory at the INS in Bogotá for confirmation and antifungal susceptibility testing (AFST). Ninety-nine *C. auris* isolates were received at the U.S. Centers for Disease Control and Prevention (CDC) with specimen collection dates between June 2016 – January 2021. Specimens were collected by convenience sampling in 41 healthcare facilities in eleven Colombian departments and they are not representative of the prevalence of *C. auris* cases in Colombia. Matrix-assisted laser desorption ionization-time of flight (MALDI-TOF) mass spectrometry was used according to the Bruker Daltonics’ protocol with minor modifications as reported previously [21] for species confirmation.

### Culture and antifungal susceptibility testing (AFST)

*C. auris* isolates were screened for elevated MICs to itraconazole, voriconazole, and fluconazole using the reference broth microdilution method as described by Clinical and Laboratory Standards Institute (CLSI) document M38[22] and for elevated MICs to amphotericin B utilizing yeast Etest strips, according to the manufacturer’s instructions (bioMerieux, Marcy l’Etoile, France). Tentative MIC breakpoints for antifungal resistance in *C. auris* have been established by CDC (https://www.cdc.gov/fungal/candida-auris/c-auris-antifungal.html). For amphotericin B, isolates with a MIC value ≥ 2 were considered resistant; if using Etest for amphotericin B and a MIC of 1.5 was determined, that value was rounded up to 2. For fluconazole, isolates with a MIC value ≥ 32 were considered resistant. For echinocandins, isolates with either caspofungin MIC value ≥ 2, anidulafungin MIC value ≥ 4 or micafungin MIC value ≥ 4 were considered resistant.

### DNA extraction and genomic sequencing

DNA was extracted using the DNeasy Blood and Tissue kit (Qiagen, Gaithersburg, MD, USA) according to the manufacturer’s recommendations. Genomic libraries were constructed using NEBNext Ultra DNA Library Prep kit (New England Biolabs, Ipswich, MA, USA) for Illumina and sequenced on Illumina Nova-Seq using NovaSeq 6000SP reagent kit (500 Cycles). Read data was deposited into the Sequence Read Archive (SRA) database at the National Center for Biotechnology Information (NCBI), under BioProject PRJNA1003896.

### Single-nucleotide polymorphism (SNP) calling using MycoSNP

In addition to the 99 Colombian isolates sequenced in this study, we included sequences from 83 Colombian isolates that had been previously described in Escandon et al., 2019 [7], and 20 clade IV isolates from the United States, Venezuela, Israel, and Panama.

An assembly of *C. auris* B11243 (GCA_003014415.1), a clade IV isolate from Venezuela, was used as the reference for read mapping and SNP calling. This reference sequence had a length of 12.3 Mb and GC content of 45%. For the analysis, MycoSNP-nf v 1.4 workflow was used [23] (https://github.com/CDCgov/mycosnp-nf). The reference genome was masked for repeats using the nucmer command from MUMmer (v 4.0) and Bedtools (v 2.29.2). The reference genome was indexed for read alignment using the BWA index command and for variant calling using Samtools (v 1.10) faidx and Picards GATK (v 2.22.9) [http://broadinstitute.github.io/picard/]. Low-quality data, trimming, and filtering were performed using FaQCs (v 2.10). The trimmed reads were used for alignment using the BWA (v 0.7.17) MEM command. Further, the aligned BAM files from each sample were pre-processed using Samtools and Picard commands and made ready for variant calling. Genome Analysis Toolkit GATK (v 4.1.4.1) was used for variant calling using the haploid mode. GATK’s VariantFiltration tool was used to filter sites based on the filtering expression “QD < 2.0 || FS > 60.0 || MQ < 40.0”. Using a customized script, genotypes were filtered if the minimum genotype quality was <50, the percentage alternate allele was <0.80 or the depth was <10.

### Phylogenetic reconstruction

MycoSNP workflow recovered all variable positions in FASTA format from the VCF files using a Python script “vcfSnpsToFasta.py” (https://github.com/broadinstitute/broad-fungalgroup/tree/master/scripts/SNPs). For the phylogenetic analysis, 1,779 polymorphic position were concatenated, the pairwise distances and neighbor-joining tree (NJ) were calculated with the MEGA-cc program [24] using 1,000 bootstraps. For tree visualization, we used the web-based JavaScript application, Microreact. The median SNP difference were calculated using in-house R-scripts and pairwise distance matrix calculated by MEGA-X.

### SNP annotations using SnpEff

SnpEff v 5.0 was used to predict the effects of associated mutations within genes [25]. From the VCF file obtained with MycoSNP workflow, SNPs were annotated using the database available in SnpEff for *Candida auris* GCA_003014415.1. To identify mutations only in coding regions, we used the parameters -no-downstream -no-upstream -no-intergenic. Several filters were applied to the annotated VCF to identify the mutations present in some protein-encoded genes associated with resistance to azoles in some *Candida* genus (*ERG11, TAC1B, PDR1, UPC2, MRR1, ERG3*) and genes associated with high MIC values of amphotericin B (PSK76257, PSK74852), at specific positions previously reported in the literature. In-house R scripts were used to compare groups of samples with different genotypes to describe differences in MIC values.

## RESULTS

### Geographic distribution and antifungal resistance of *Candida auris* phylogenetic clusters

Ninety-nine *C. auris* isolates with specimen collection dates ranging from June 2016–January 2021 were newly sequenced in this study (Supplementary Table 1). Specimen types included blood (n=68, 69%), urine (n=9, 9%), skin swabs (n=4, 4%), oral (n=3, 3%), nasal (n=3, 3%), and other sources (n=12, 12%).

Phylogenetic analysis of the 99 newly generated sequences, and the 83 previously generated sequences revealed that all *C. auris* in Colombia were clade IV (maximum SNP difference to the reference sequence: 205 SNPs). Overall, isolates predominately clustered by country. Colombian isolates were dispersed in two clusters, referred to as clusters C1 and C2. Most Colombian isolates (n=166, 91%) collected between June 2016 – January 2021 clustered with the previously described Colombian isolates in cluster C1 with collection dates from February 2015 – August 2016. The median SNP difference within cluster C1 was 38 SNPs (range: 0 - 76 SNPs; Figure 2A).

**Figure 1:**
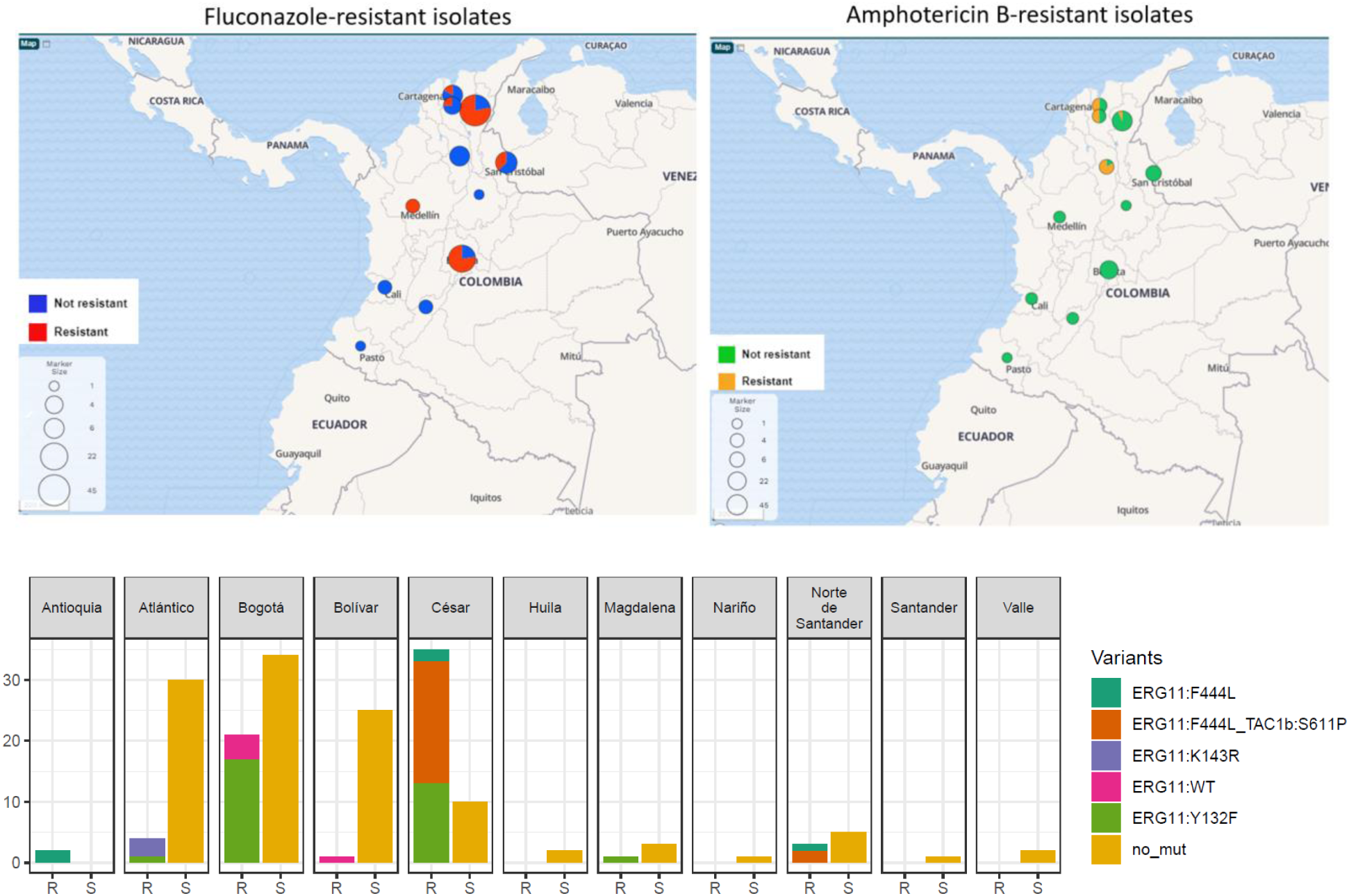
**A)** Geographic distribution of antifungal-resistant isolates. **B)** Bar plot of the number of susceptible/resistant isolates for each Colombian state. Bars corresponding to the resistant (R) isolates are divided by colors that represent the proportion of different mutations. Yellow bars represent susceptible (S) isolates. no mut=Non identified mutations in *ERG11* gene.

**Figure 2:**
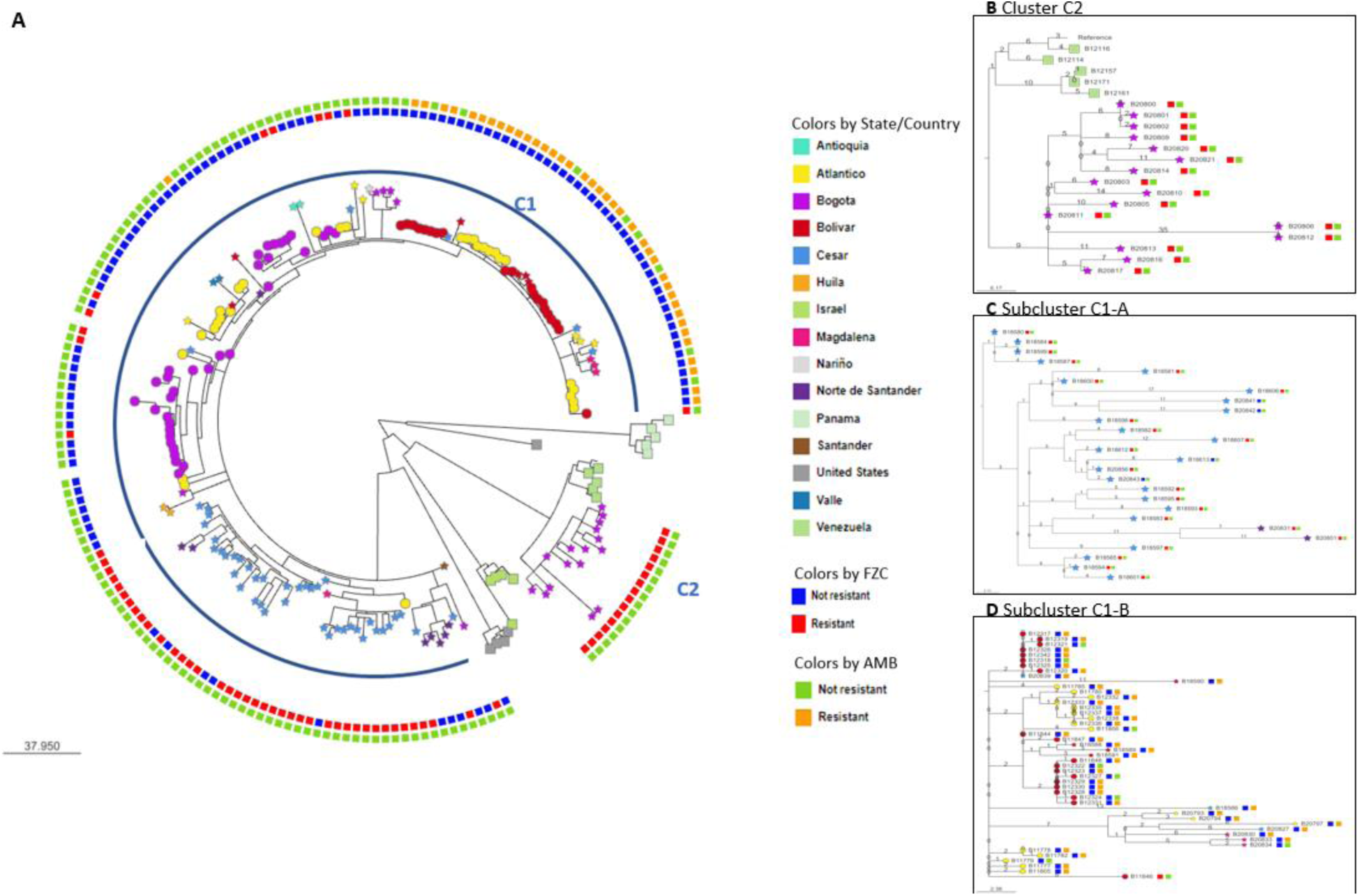
**A)** Phylogenetic tree using the Neighbor-Joining method, the majority of isolates cluster by country. Cluster C1 consist of Colombian Isolates, Cluster C2 consist of Venezuelan Isolates and fluconazole-resistant Colombian isolates. Taxa color represent Colombian state or Country, isolates from countries other than Colombia are shown as squares, Colombian isolates sequenced in previous studies are shown as circles and Colombian isolates sequenced in this study are shown as stars. External circles colors represent resistant and no resistant phenotypes for fluconazole and amphotericin B. **B)** Zoom in view of Cluster C2. **C)** Zoom in view of subcluster A, which contains manly fluconazole-resistant isolates **D)** Zoom in view of subcluster B, which contains manly amphotericin B-resistant isolates. For panels B, C and D; the numbers on the branches represent the number of SNPs between isolates and the squares to the right of the label represent resistant and non-resistant phenotypes.

Cluster C2 was separated by 152 SNPs from cluster C1. Cluster C2 comprised 16 isolates from Bogotá collected between May - November 2020 and five isolates from Venezuela collected between October 2012- May 2015 (Figure 2A). All Colombian isolates in this cluster were fluconazole-resistant, as well as two Venezuelan isolates (supplementary table 1). The median SNP difference within cluster C2 was 23 SNPs (range: 0 - 55 SNPs; Figure 2B). All 16 C2 Colombian isolates were collected in 11 different facilities in Bogotá, including a pair of isolates (B20806 and B20812) from two different facilities that were 0 SNPs apart.

Within cluster C1, we highlighted two subclusters of isolates with bootstrap values of at least 98% and with high (>80%) proportion of antifungal-resistant isolates (supplementary figure 2). These subclusters were C1-A and C1-B with 26 and 47 isolates, respectively, and the remaining 93 isolates in C1 were susceptible or fluconazole-resistant (Figure 2A). The subcluster C1-A, contained isolates mainly from César and the median SNP difference was 19 SNPs (range: 0 - 45 SNPs; Figure 2C). Of the 26 isolates in C1-A, 22 (85%) were resistant to fluconazole. The subcluster C1-B, comprised isolates from the north coast (Bolívar, Atlántico, César, and Magdalena) and the median SNP difference was 9 SNPs (range: 0 - 31 SNPs; Figure 2D). Of these 47 isolates in C1-B, 38 (81%) isolates were resistant to amphotericin B.

### Molecular characterization of antifungal-resistant isolates

Among all 182 Colombian isolates; 67 (37%) were resistant to fluconazole, 39 (21%) were resistant to amphotericin B, and only one was resistant to anidulafungin. However, three identified as susceptible to anidulafungin displayed high MIC values of 2 (Supplemental Figure 1). We found that 62 (93%) of the 67 fluconazole-resistant isolates had known mutations in *ERG11* and *TAC1B* genes. The Y132F mutation in the *ERG11* gene was the most prevalent mutation, observed in 32 (48%) azole-resistant isolates (Supplementary table 2). Isolates with mutation Y132F in the *ERG11* gene had significantly higher fluconazole MIC values (MIC value median = 256) than isolates with concomitant mutations F444L in the *ERG11* gene and S611P in *TAC1B* gene (MIC value median = 64). The isolates with concomitant mutations also had significantly higher fluconazole MIC values than isolates with the single mutation F444L in the *ERG11* gene (MIC value median = 32) (Figure 3). All fluconazole-resistant isolates in subcluster C1-A (n=22), collected in 2016, carried the concomitant mutations F444L in the *ERG11* gene and S611P in *TAC1B* gene, and all (n= 16) fluconazole-resistant isolates in clade C2 carried the mutation Y132.

**Figure 3:**
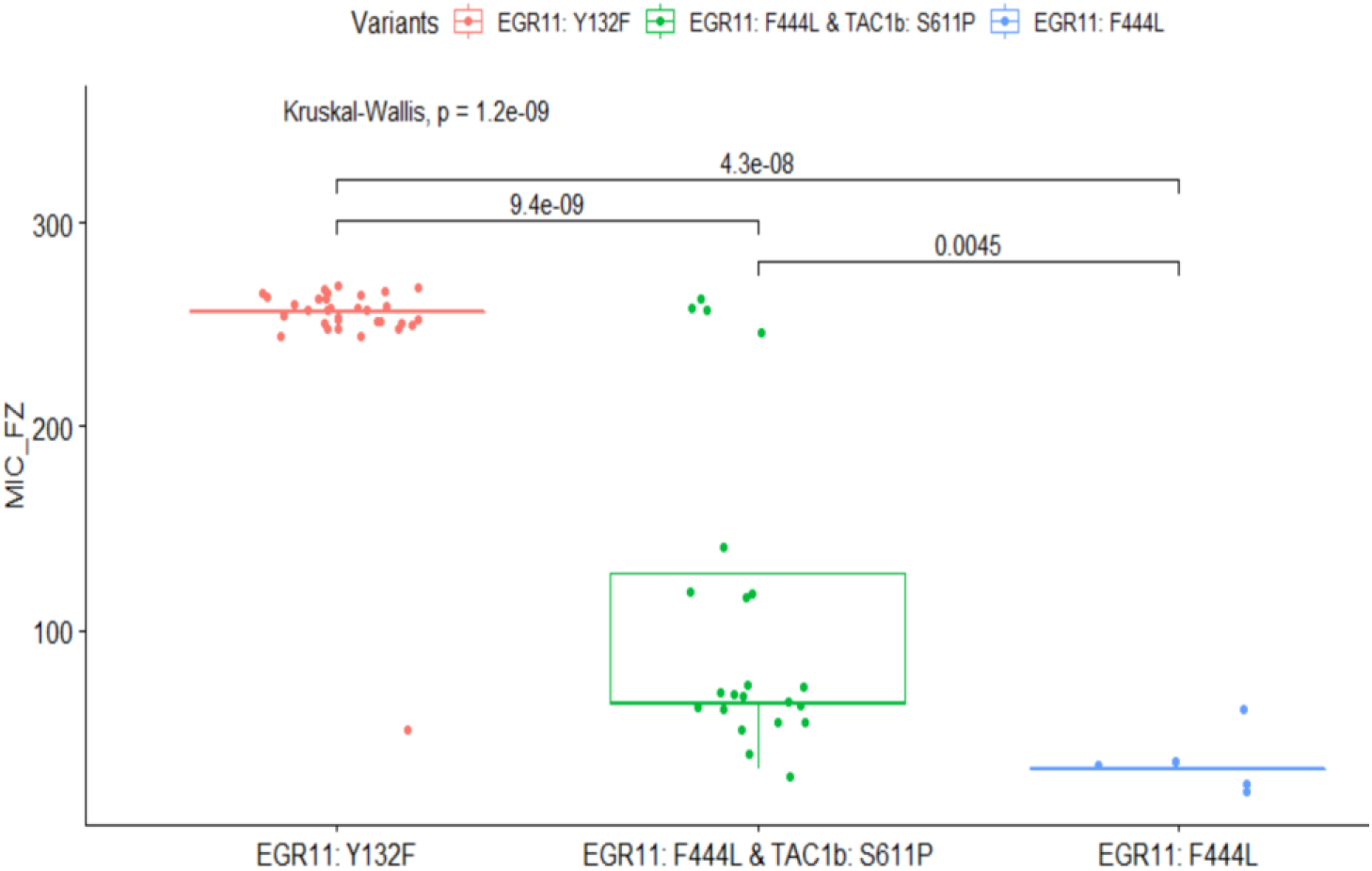
R boxplot on common mutation in *ERG11* and *TAC1B* genes and MIC values (mg/µL) of fluconazole-resistant isolates. Significant differences between MIC values support the cumulative effect of the concomitant mutation in ERG11 and *TAC1B* genes. Isolates B11790 and B11087 with mutations *ERG11* K143R was excluded from the graphic. Jitter R function was used to add minimal and random noise to the MIC values to visualize the number of isolates at each value.

Of the 39 amphotericin B-resistant isolates, we found that 38 (97%) carried the substitutions S108N in *FLO8* (PSK76257) gene and I139T PSK74852 gene. Both substitutions were also found in eight susceptible isolates with amphotericin B MIC values ≥ 0.75 and in only one isolate with amphotericin B MIC value 0.25 mg/µL.

## DISCUSSION

In this study, we provided a comprehensive update on the genomic epidemiology of *C. auris* in Colombia. One of the main findings was that all *C. auris* cases in Colombia continue to be of clade IV. In contrast to several countries experiencing outbreaks and reporting *C auris* cases over multiple years have identified isolates from more than one major *C. auris* clade in their country [10]. Additionally, Clade I has been reported in nearby countries in Latin America [26]. The uniformity of clade IV isolates could be due to 1) lack of introductions of other clades or 2) lack of subsequent transmission after introductions of other clades. Alternatively, isolates from other *C. auris* clades could have been missed by our collection efforts.

Previously, a global molecular epidemiologic description of *C. auris* reported genetic clustering by country within clade IV [10]. Here, we observed a weaker phylogeographic structure as isolates from Colombia were dispersed into two clusters, one also containing isolates from Venezuela. Specifically, this cluster C2 included fluconazole-resistant isolates collected in 2020 from Bogotá clustered with isolates from cases in Venezuela, and the remaining Colombian isolates clustered with previously described isolates from cases in Colombia (Cluster C1). Interestingly, all clinical isolates from Bogotá collected before 2020 were susceptible; this could suggest a later introduction of fluconazole-resistant resistant cases from Venezuela. Given the observed phylogeny, we hypothesize that an introduction of clade IV cases occurred in Colombia, possibly from Venezuela where *C. auris* is circulating since 2012. Such a recent introduction would have (in early 2020) given rise to the “C2” cluster described above. Additional studies of phylogeographic and molecular clock analyses are needed to better estimate when this described introduction could have occurred.

Additionally, low median SNP differences were observed among isolates in both subclusters C1- A and C1-B spanning a six-year time period, suggesting that most cases in Colombia (cluster C1) are the result of ongoing transmission since 2016.

By pairing epidemiologic, genomic, and AFST data, we were able to conclude that amphotericin B-resistant isolates clustered (subcluster C1-B) and were predominately collected from the northern region of the country, which was previously reported by Escandon et al. in 2019 [7]. Additionally, the fluconazole-resistant isolates from Colombia were predominately collected from the northeast region and the capital city, Bogotá, in the central region.

Regarding the genotypes associated with fluconazole resistance, we observed that the most common mutation was *ERG11* Y132F, consistent with Y132F being previously reported as the most prevalent mutation in fluconazole-resistant clade IV isolates [10]. In Colombia, Y132F was also associated with the highest MICs for fluconazole compared to other mutations (Figure 3). This mutation was present in all Colombian isolates in Cluster C2 from 2020. The second most common genotype associated with fluconazole resistance had both mutations in *ERG11* (F444L) and *TAC1B* (S611P; Figure 3). Interestingly, the genotype *ERG11* F444L and *TAC1B* S611P has been reported only in Colombian clade IV isolates to date [12] and it appeared restricted to isolates from subcluster C1-A, which were collected in 2016 or later years. The emergence of a new mechanism of fluconazole resistance in 2016 as well as the introduction of resistant *C. auris* from another country in 2020, could explain the observed increase in MIC values of fluconazole-resistant isolates over time. A similar finding was first reported by Escandon et. al 2022 [8], who proposed the transmission of fluconazole-resistant isolates as a possible factor responsible for the increase of fluconazole resistance in the country. Further studies on serial isolates of *C. auris* from the same patient would be needed to investigate if the observed increase of fluconazole resistance is also related with acquired resistance in response to treatment.

Notably, we did not identify mutations in either *ERG11* or *TAC1B* genes in the five azole-resistant isolates; these isolates also did not have mutations in *PDR1, UPC2, MRR1,* and *ERG3* genes. Future work may investigate other potential mechanisms of azole resistance including copy number variants (CNVs) and overexpression of genes.

In contrast with the observed increase in MIC values of fluconazole-resistant isolates, we found that resistance to amphotericin B has not changed significantly over time in Colombia. In amphotericin B-resistant isolates, we spotted the substitution S108N in *FLO8* (PSK76257) and the substitution I139T in PSK74852, which were also previously reported in Colombian isolates by Escandon et al 2019. If and how the described substitutions lead to a reduction in amphotericin B susceptibility is unknown.

A limitation of this work is that clinical information including history of antifungal treatment and patients’ treatment outcomes was unknown. Some resistant isolates could have acquired resistance in response to therapy rather than transmission of drug-resistant *C. auris*. However, given the low genetic differences between some cases and the common phylogeny, transmission of fluconazole-resistant *C. auris* seems more likely. A second limitation is that specimens were collected by convenience sampling and therefore are not representative of *C. auris* cases in Colombia.

In conclusion, the findings of this work provided evidence of ongoing spread and a better understanding of molecular mechanisms associated with *C. auris* antifungal resistance in Colombia. These findings contributed to the understanding of the *C. auris* epidemiology and factors associated with transmission in Colombia and could be useful to guide prevention and control strategies. Additionally, these findings help to identify mechanisms conferring fluconazole and amphotericin B resistance in Colombia.

## ACKNOWLEDGMENTS

We give special thanks to Anastasia P. Litvintseva for the insightful discussions and CDC’s Office of Advanced Molecular Detection (OAMD). FINANCIAL SUPPORT We acknowledge Oak Ridge Institute for Science and Education (ORISE) for financial support for EM.

## DECLARATION OF INTERESTS

The authors report no conflicts of interest.

## DISCLAIMER

The findings and the conclusions in this report are those of the authors and do not necessarily represent the views of the Centers for Disease Control and Prevention. Use of trade names is for identification only and does not imply endorsement.

---

**Supplemental Figure 1:**
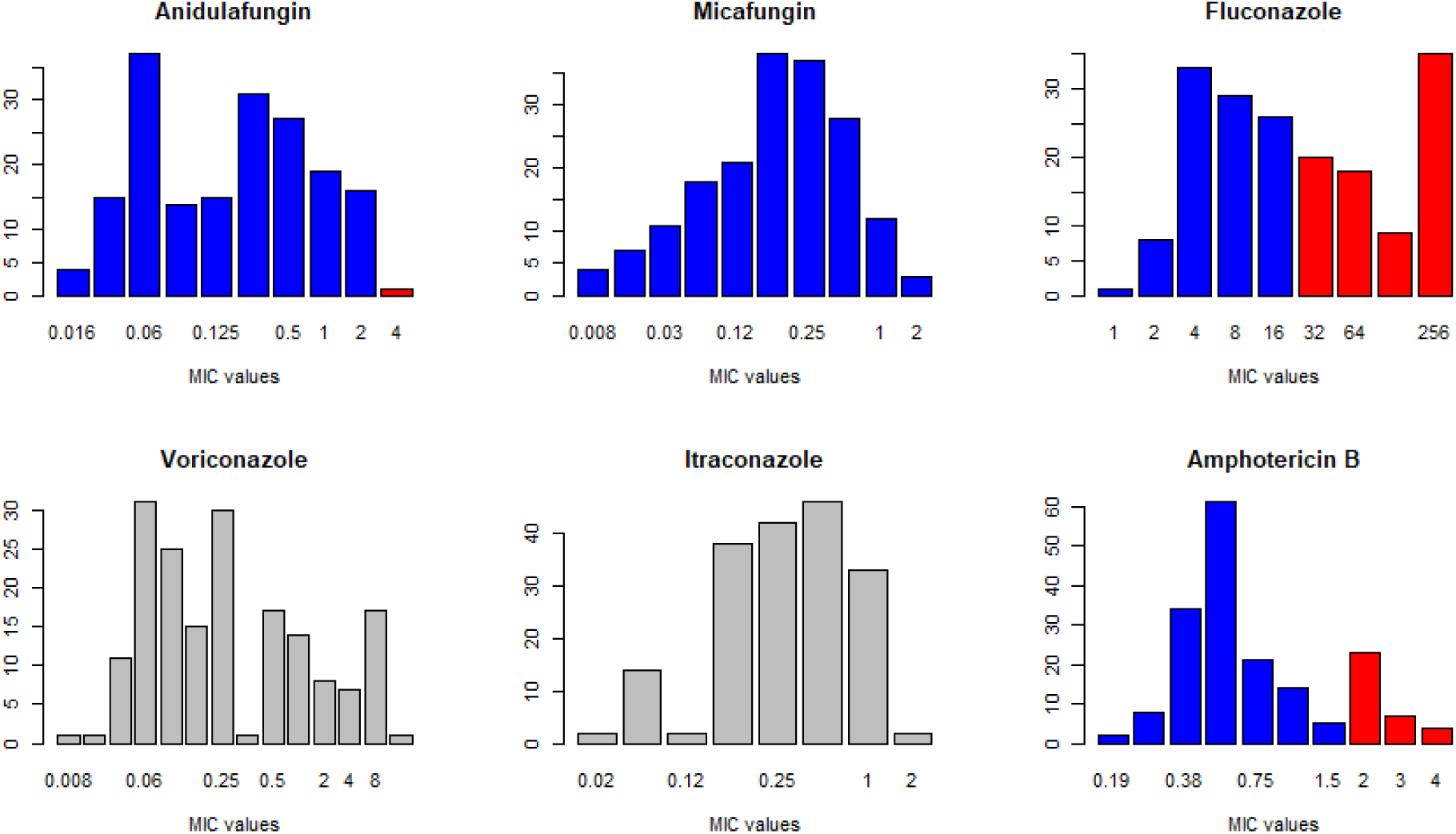
MIC value frequency distribution. Antifungal susceptibility testing (AFST). Red and blue bars indicate resistant and susceptible isolates respectively, according to tentative MIC breakpoints. Grey bars represent that there are not tentative MIC breakpoints for the specific drug.

**Supplemental Figure 2:**
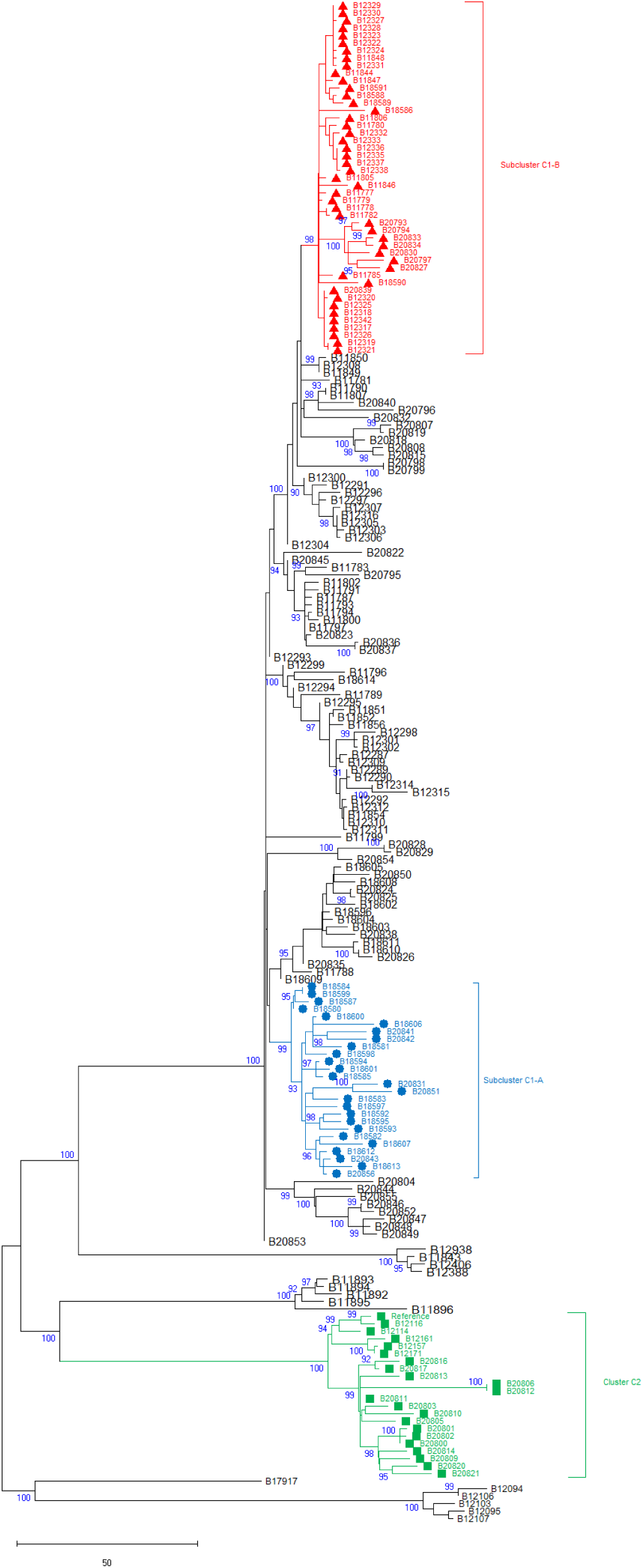
NJ tree, bootstrap 1000, bootstrap values lower than 90 were hidden.

**Supplementary table 1:** Metadata and accession numbers for *C. auris* isolates.

| Bnumber | Specimen collected date | Specimen source Type | Country | Colombian departament | AND MIC | CAS MIC | MF MIC | AMB MIC | FZ MIC | Variants | SRR | Citation |
| --- | --- | --- | --- | --- | --- | --- | --- | --- | --- | --- | --- | --- |
| B11777 | 2/3/2015 | Blood | Colombia | Atlántico | 2 | NA | 0.5 | 2 | 4 | NA | SRR7140008 | Escandon et al. 2019 |
| B11778 |  | Blood | Colombia | Atlántico | 4 | NA | 0.5 | 3 | 8 | NA | SRR7140036 | Escandon et al. 2019 |
| B11779 |  | Blood | Colombia | Atlántico | 2 | NA | 0.5 | 1 | 4 | NA | SRR7140057 | Escandon et al. 2019 |
| B11780 | 9/10/2015 | Blood | Colombia | Atlántico | 2 | NA | 0.5 | 3 | 4 | NA | SRR7140031 | Escandon et al. 2019 |
| B11781 | 3/16/2016 | Blood | Colombia | Atlántico | 2 | NA | 0.5 | 0.38 | 4 | NA | SRR7140033 | Escandon et al. 2019 |
| B11782 | 11/25/2015 | Blood | Colombia | Atlántico | 2 | NA | 0.5 | 3 | 4 | NA | SRR7140067 | Escandon et al. 2019 |
| B11783 |  | Blood | Colombia | Atlántico | 2 | NA | 0.5 | 0.5 | 8 | NA | SRR7140056 | Escandon et al. 2019 |
| B11785 |  | Blood | Colombia | Atlántico | 2 | NA | 0.5 | 2 | 8 | NA | SRR7140021 | Escandon et al. 2019 |
| B11787 | 5/13/2015 | Blood | Colombia | Atlántico | 2 | NA | 0.5 | 0.38 | 16 | NA | SRR7140012 | Escandon et al. 2019 |
| B11788 | 8/15/2015 | Blood | Colombia | Atlántico | 2 | NA | 0.5 | 0.38 | 64 | EGR11: Y132F | SRR7140041 | Escandon et al. 2019 |
| B11789 |  | Blood | Colombia | Atlántico | 2 | NA | 1 | 0.5 | 16 | NA | SRR7140082 | Escandon et al. 2019 |
| B11790 |  | Blood | Colombia | Atlántico | 1 | NA | 0.5 | 0.5 | 32 | EGR11: K143R | SRR7140029 | Escandon et al. 2019 |
| B11791 | 9/7/2015 | Blood | Colombia | Atlántico | 0.5 | NA | 0.25 | 0.5 | 16 | NA | SRR7140070 | Escandon et al. 2019 |
| B11793 | 7/15/2015 | Blood | Colombia | Atlántico | 0.5 | NA | 0.25 | 0.25 | 8 | NA | SRR7140071 | Escandon et al. 2019 |
| B11794 | 7/2/2015 | Blood | Colombia | Atlántico | 1 | NA | 0.5 | 0.38 | 16 | NA | SRR7140055 | Escandon et al. 2019 |
| B11796 | 3/12/2016 | Blood | Colombia | Atlántico | NA | NA | NA | NA | NA | NA | SRR10461264 | Chow et al. 2020 |
| B11797 |  | Blood | Colombia | Atlántico | 1 | NA | 0.5 | 0.38 | 8 | NA | SRR7140015 | Escandon et al. 2019 |
| B11799 |  | Blood | Colombia | Atlántico | 2 | NA | 2 | 0.5 | 16 | NA | SRR7140004 | Escandon et al. 2019 |
| B11800 |  | Blood | Colombia | Atlántico | 0.5 | NA | 0.5 | 0.38 | 16 | NA | SRR7140002 | Escandon et al. 2019 |
| B11802 |  | Blood | Colombia | Atlántico | 0.25 | NA | 0.5 | 0.25 | 16 | NA | SRR7140077 | Escandon et al. 2019 |
| B11805 | 4/13/2015 | Blood | Colombia | Atlántico | 1 | NA | 0.5 | 2 | 4 | NA | SRR7140075 | Escandon et al. 2019 |
| B11806 |  | Blood | Colombia | Atlántico | 1 | NA | 0.5 | 1 | 4 | NA | SRR7140016 | Escandon et al. 2019 |
| B11807 | 1/3/2015 | Blood | Colombia | Atlántico | 1 | NA | 0.5 | 0.38 | 32 | EGR11: K143R | SRR7140080 | Escandon et al. 2019 |
| B11843 | 5/21/2016 | Blood | United States | NA | NA | NA | NA | NA | NA | NA | SRR7909220 | Chow et al. 2020 |
| B11844 | 7/15/2016 | Blood | Colombia | Bolivar | 1 | NA | 1 | 2 | 4 | NA | SRR7140078 | Escandon et al. 2019 |
| B11846 |  | Blood | Colombia | Bolivar | 0.25 | NA | 0.5 | 0.25 | 32 | EGR11: WT | SRR7140064 | Escandon et al. 2019 |
| B11847 |  | Blood | Colombia | Bolivar | 1 | NA | 1 | 4 | 4 | NA | SRR7140066 | Escandon et al. 2019 |
| B11848 |  | Blood | Colombia | Bolivar | 2 | NA | 2 | 3 | 4 | NA | SRR7140065 | Escandon et al. 2019 |
| B11849 |  | Blood | Colombia | Bogotá | 1 | NA | 0.5 | 0.38 | 4 | NA | SRR7140045 | Escandon et al. 2019 |
| B11850 |  | Blood | Colombia | Bogotá | 0.5 | NA | 1 | 0.5 | 8 | NA | SRR7140039 | Escandon et al. 2019 |
| B11851 |  | Blood | Colombia | Bogotá | 1 | NA | 1 | 0.38 | 32 | NA | SRR7140040 | Escandon et al. 2019 |
| B11852 | 6/15/2016 | Blood | Colombia | Bogotá | 1 | NA | 1 | 0.5 | 32 | NA | SRR7140009 | Escandon et al. 2019 |
| B11854 | 7/28/2016 | urine | Colombia | Bogotá | NA | NA | NA | NA | NA | NA | SRR10461261 | Chow et al. 2020 |
| B11856 |  | Blood | Colombia | Bogotá | 2 | NA | 1 | 0.5 | 4 | NA | SRR7140054 | Escandon et al. 2019 |
| B11892 | 10/15/2014 | Blood | Israel | NA | NA | NA | NA | NA | NA | NA | SRR10461259 | Chow et al. 2020 |
| B11893 | 5/15/2014 | Blood | Israel | NA | NA | NA | NA | NA | NA | NA | SRR10461258 | Chow et al. 2020 |
| B11894 | 8/15/2014 | Blood | Israel | NA | NA | NA | NA | NA | NA | NA | SRR10461257 | Chow et al. 2020 |
| B11895 | 5/15/2014 | Blood | Israel | NA | NA | NA | NA | NA | NA | NA | SRR10461256 | Chow et al. 2020 |
| B11896 | 4/15/2015 | Urine | Israel | NA | NA | NA | NA | NA | NA | NA | SRR10461255 | Chow et al. 2020 |
| B12094 | 9/22/2016 | Urine | Panamá | NA | NA | NA | NA | NA | NA | NA | SRR10461252 | Chow et al. 2020 |
| B12095 | 8/2/2016 | Urine | Panamá | NA | NA | NA | NA | NA | NA | NA | SRR10461251 | Chow et al. 2020 |
| B12103 | 7/25/2016 | Urine | Panamá | NA | NA | NA | NA | NA | NA | NA | SRR10461181 | Chow et al. 2020 |
| B12106 | 9/21/2016 | Urine | Panamá | NA | NA | NA | NA | NA | NA | NA | SRR10461179 | Chow et al. 2020 |
| B12107 | 9/19/2016 | Urine | Panamá | NA | NA | NA | NA | NA | NA | NA | SRR10461178 | Chow et al. 2020 |
| B12114 | 10/16/2012 | Blood | Venezuela | NA | NA | 0.25 | NA | 0.5 | 256 | NA | SRR10461177 | Chow et al. 2020 |
| B12116 | 10/20/2012 | Blood | Venezuela | NA | NA | 0.094 | NA | 0.008 | 256 | NA | SRR10461176 | Chow et al. 2020 |
| B12157 | 1/9/2015 | CSF | Venezuela | NA | NA | 4 | NA | 0.5 | 16 | NA | SRR10461126 | Chow et al. 2020 |
| B12161 | 1/5/2015 | Blood | Venezuela | NA | NA | NA | NA | NA | NA | NA | SRR10461124 | Chow et al. 2020 |
| B12171 | 5/30/2015 | CSF | Venezuela | NA | NA | NA | NA | NA | NA | NA | SRR10461206 | Chow et al. 2020 |
| B12287 |  | Floor | Colombia | Bogotá | 0.06 | NA | 0.25 | 0.5 | 16 | NA | SRR7140047 | Escandon et al. 2019 |
| B12289 |  | Floor | Colombia | Bogotá | 0.125 | NA | 0.25 | 0.5 | 16 | NA | SRR7140059 | Escandon et al. 2019 |
| B12290 |  | Bed rail | Colombia | Bogotá | 0.06 | NA | 0.125 | 0.5 | 16 | NA | SRR7140042 | Escandon et al. 2019 |
| B12291 |  | Night table | Colombia | Bogotá | 0.25 | NA | 0.25 | 0.5 | 8 | NA | SRR7140017 | Escandon et al. 2019 |
| B12292 |  | Closet | Colombia | Bogotá | 0.25 | NA | 0.25 | 0.5 | 32 | EGR11: WT | SRR7140058 | Escandon et al. 2019 |
| B12293 |  | Towel | Colombia | Bogotá | 0.25 | NA | 0.25 | 0.5 | 64 | EGR11: WT | SRR7140032 | Escandon et al. 2019 |
| B12294 |  | Cell phone | Colombia | Bogotá | 0.125 | NA | 0.25 | 0.5 | 32 | EGR11: WT | SRR7140006 | Escandon et al. 2019 |
| B12295 |  | Bedpan | Colombia | Bogotá | 0.06 | NA | 0.25 | 0.5 | 32 | EGR11: WT | SRR7140025 | Escandon et al. 2019 |
| B12296 |  | Mattress | Colombia | Bogotá | 0.125 | NA | 0.125 | 0.5 | 4 | NA | SRR7140038 | Escandon et al. 2019 |
| B12297 |  | Chair. towel | Colombia | Bogotá | 0.125 | NA | 0.25 | 0.5 | 8 | NA | SRR7140074 | Escandon et al. 2019 |
| B12298 |  | Bedpan | Colombia | Bogotá | 0.06 | NA | 0.25 | 0.5 | 16 | NA | SRR7140043 | Escandon et al. 2019 |
| B12299 |  | Mop | Colombia | Bogotá | 0.06 | NA | 0.125 | 0.38 | 16 | NA | SRR7140010 | Escandon et al. 2019 |
| B12300 |  | TV control | Colombia | Bogotá | 0.06 | NA | 0.125 | 0.5 | 16 | NA | SRR7140069 | Escandon et al. 2019 |
| B12301 |  | Meal table | Colombia | Bogotá | 0.125 | NA | 0.25 | 0.38 | 8 | NA | SRR7140001 | Escandon et al. 2019 |
| B12302 |  | Chair | Colombia | Bogotá | NA | NA | NA | NA | NA | NA | SRR7140003 | Escandon et al. 2019 |
| B12303 |  | Sink | Colombia | Bogotá | 0.06 | NA | 0.125 | 0.5 | 8 | NA | SRR7140030 | Escandon et al. 2019 |
| B12304 |  | Toilet | Colombia | Bogotá | 0.06 | NA | 0.125 | 0.38 | 8 | NA | SRR7140044 | Escandon et al. 2019 |
| B12305 |  | Wheelchair | Colombia | Bogotá | 0.06 | NA | 0.125 | 0.38 | 8 | NA | SRR7140022 | Escandon et al. 2019 |
| B12306 |  | Closet | Colombia | Bogotá | 0.06 | NA | 0.125 | 0.5 | 8 | NA | SRR7140013 | Escandon et al. 2019 |
| B12307 |  | Mop bucket | Colombia | Bogotá | 0.06 | NA | 0.125 | 0.5 | 8 | NA | SRR7140052 | Escandon et al. 2019 |
| B12308 |  | Floor | Colombia | Bogotá | 0.125 | NA | 0.125 | 0.38 | 4 | NA | SRR7140061 | Escandon et al. 2019 |
| B12309 |  | Nurse shoes | Colombia | Bogotá | 0.06 | NA | 0.125 | 0.5 | 16 | NA | SRR7140024 | Escandon et al. 2019 |
| B12310 |  | Sink | Colombia | Bogotá | 0.06 | NA | 0.25 | 0.5 | 8 | NA | SRR7140007 | Escandon et al. 2019 |
| B12311 |  | Door | Colombia | Bogotá | 0.125 | NA | 0.25 | 0.5 | 16 | NA | SRR7140020 | Escandon et al. 2019 |
| B12312 |  | Bedpan | Colombia | Bogotá | 0.125 | NA | 0.125 | 0.38 | 16 | NA | SRR7140068 | Escandon et al. 2019 |
| B12314 |  | Left groin | Colombia | Bogotá | 0.125 | NA | 0.125 | 0.5 | 16 | NA | SRR7140063 | Escandon et al. 2019 |
| B12315 |  | Rectum | Colombia | Bogotá | 0.06 | NA | 0.125 | 0.5 | 8 | NA | SRR7140076 | Escandon et al. 2019 |
| B12316 |  | Right groin | Colombia | Bogotá | 0.06 | NA | 0.06 | 0.38 | 8 | NA | SRR7140035 | Escandon et al. 2019 |
| B12317 |  | Floor | Colombia | Bolivar | 0.06 | NA | 0.125 | 3 | 4 | NA | SRR7140005 | Escandon et al. 2019 |
| B12318 |  | Equipment | Colombia | Bolivar | 0.03 | NA | 0.125 | 1 | 4 | NA | SRR7140053 | Escandon et al. 2019 |
| B12319 |  | Body cables | Colombia | Bolivar | 0.06 | NA | 0.125 | 2 | 2 | NA | SRR7140081 | Escandon et al. 2019 |
| B12320 |  | Bed | Colombia | Bolivar | 0.06 | NA | 0.125 | 2 | 2 | NA | SRR7140079 | Escandon et al. 2019 |
| B12321 |  | Equipment | Colombia | Bolivar | 0.06 | NA | 0.25 | 0.75 | 2 | NA | SRR7140060 | Escandon et al. 2019 |
| B12322 |  | Bed rail | Colombia | Bolivar | 0.06 | NA | 0.125 | 1 | 4 | NA | SRR7140072 | Escandon et al. 2019 |
| B12323 |  | Mattress | Colombia | Bolivar | 0.06 | NA | 0.06 | 2 | 4 | NA | SRR7140014 | Escandon et al. 2019 |
| B12324 |  | Body cables | Colombia | Bolivar | 0.06 | NA | 0.03 | 1 | 4 | NA | SRR7140049 | Escandon et al. 2019 |
| B12325 |  | Bed rails | Colombia | Bolivar | 0.06 | NA | 0.125 | 2 | 4 | NA | SRR7140046 | Escandon et al. 2019 |
| B12326 |  | Stretcher | Colombia | Bolivar | 0.06 | NA | 0.125 | 2 | 4 | NA | SRR7140050 | Escandon et al. 2019 |
| B12327 |  | Left armpit | Colombia | Bolivar | 0.016 | NA | 0.06 | 0.75 | 2 | NA | SRR7140023 | Escandon et al. 2019 |
| B12328 |  | Right armpit | Colombia | Bolivar | 0.03 | NA | 0.125 | 2 | 4 | NA | SRR7140027 | Escandon et al. 2019 |
| B12329 |  | Groin composite | Colombia | Bolivar | 0.03 | NA | 0.125 | 2 | 2 | NA | SRR7140037 | Escandon et al. 2019 |
| B12330 |  | Hands | Colombia | Bolivar | 0.06 | NA | 0.125 | 2 | 4 | NA | SRR7140073 | Escandon et al. 2019 |
| B12331 |  | Hands | Colombia | Bolivar | 0.03 | NA | 0.125 | 2 | 2 | NA | SRR7140026 | Escandon et al. 2019 |
| B12332 |  | Right ear | Colombia | Atlántico | 0.03 | NA | 0.125 | 2 | 1 | NA | SRR7140019 | Escandon et al. 2019 |
| B12333 |  | Right armpit | Colombia | Atlántico | 0.03 | NA | 0.125 | 2 | 2 | NA | SRR7140034 | Escandon et al. 2019 |
| B12335 |  | Left ear | Colombia | Atlántico | 0.06 | NA | 0.125 | 2 | 4 | NA | SRR7140048 | Escandon et al. 2019 |
| B12336 |  | Left nostril | Colombia | Atlántico | 0.06 | NA | 0.125 | 2 | 4 | NA | SRR7140028 | Escandon et al. 2019 |
| B12337 |  | Left armpit | Colombia | Atlántico | 0.06 | NA | 0.125 | 3 | 4 | NA | SRR7140051 | Escandon et al. 2019 |
| B12338 |  | Rectum | Colombia | Atlántico | 0.06 | NA | 0.125 | 2 | 4 | NA | SRR7140011 | Escandon et al. 2019 |
| B12342 |  | Stethoscope | Colombia | Bolivar | 0.06 | NA | 0.125 | 4 | 4 | NA | SRR13292041 | Escandon et al. 2019 |
| B12388 | 9/25/2016 | Axilla/Groin swab | United States | NA | NA | NA | NA | NA | NA | NA | SRR7909221 | Chow et al. 2020 |
| B12406 | 12/27/2016 | Urine | United States | NA | NA | NA | NA | NA | NA | NA | SRR7909405 | Chow et al. 2020 |
| B12938 | 4/17/2017 | Axilla/Groin swab | United States | NA | NA | NA | NA | NA | NA | NA | SRR7909357 | Chow et al. 2020 |
| B17917 | 11/30/2018 | Toenail | United States | NA | NA | NA | NA | NA | NA | NA | NA | unpublished |
| B18580 | 2/8/2017 | Blood | Colombia | Cesar | 0.016 | 0.016 | 0.03 | 0.75 | 128 | EGR11: F444L & TAC1b: S611P | SRR25583648 | This study |
| B18581 | 2/8/2017 | Blood | Colombia | Cesar | 0.06 | 0.06 | 0.016 | 1 | 64 | EGR11: F444L & TAC1b: S611P | SRR25583647 | This study |
| B18582 | 2/8/2017 | Blood | Colombia | Cesar | 0.016 | 0.016 | 0.03 | 0.75 | 128 | EGR11: F444L & TAC1b: S611P | SRR25583636 | This study |
| B18583 | 2/8/2017 | Blood | Colombia | Cesar | 0.06 | 0.06 | 0.03 | 0.5 | 64 | EGR11: F444L & TAC1b: S611P | SRR25583625 | This study |
| B18584 | 2/8/2017 | Blood | Colombia | Cesar | 0.03 | 0.03 | 0.016 | 0.5 | 64 | EGR11: F444L & TAC1b: S611P | SRR25583582 | This study |
| B18585 | 2/8/2017 | Blood | Colombia | Cesar | 0.016 | 0.016 | 0.016 | 0.75 | 128 | EGR11: F444L & TAC1b: S611P | SRR25583571 | This study |
| B18586 | 2/15/2018 | Oral swab | Colombia | Cesar | 0.125 | 0.06 | 0.06 | 1.5 | 2 | NA | SRR25583620 | This study |
| B18587 | 1/20/2017 | Blood | Colombia | Cesar | 0.03 | 0.03 | 0.016 | 0.5 | 64 | EGR11: F444L & TAC1b: S611P | SRR25583609 | This study |
| B18588 | 1/20/2017 | Blood | Colombia | Bolivar | 0.125 | 0.06 | 0.125 | 1.5 | 4 | NA | SRR25583598 | This study |
| B18589 | 2/4/2017 | Blood | Colombia | Bolivar | 0.25 | 0.125 | 0.25 | 2 | 4 | NA | SRR25583559 | This study |
| B18590 | 2/24/2017 | Blood | Colombia | Bolivar | 0.125 | 0.125 | 0.25 | 4 | 4 | NA | SRR25583646 | This study |
| B18591 | 1/13/2018 | Blood | Colombia | Bolivar | 0.5 | 0.125 | 0.125 | 2 | 4 | NA | SRR25583645 | This study |
| B18592 | 2/8/2017 | Blood | Colombia | Cesar | 0.125 | 0.06 | <0.008 | 0.5 | 64 | EGR11: F444L & TAC1b: S611P | SRR25583644 | This study |
| B18593 | 6/11/2016 | Blood | Colombia | Cesar | 0.25 | 0.06 | 0.016 | 0.75 | 64 | EGR11: F444L & TAC1b: S611P | SRR25583643 | This study |
| B18594 | 2/8/2017 | Blood | Colombia | Cesar | 0.03 | 0.03 | <0.008 | 0.5 | 32 | EGR11: F444L & TAC1b: S611P | SRR25583642 | This study |
| B18595 | 7/8/2016 | Blood | Colombia | Cesar | 0.25 | 0.125 | 0.016 | 0.5 | 32 | EGR11: F444L & TAC1b: S611P | SRR25583641 | This study |
| B18596 | 9/13/2016 | Blood | Colombia | Cesar | 0.25 | 0.06 | 0.06 | 0.38 | 256 | EGR11: Y132F | SRR25583640 | This study |
| B18597 | 7/29/2016 | Blood | Colombia | Cesar | 0.03 | 0.016 | <0.008 | 0.25 | 64 | EGR11: F444L & TAC1b: S611P | SRR25583639 | This study |
| B18598 | 7/31/2016 | Blood | Colombia | Cesar | 0.06 | 0.03 | 0.03 | 0.5 | 64 | EGR11: F444L & TAC1b: S611P | SRR25583638 | This study |
| B18599 | 2/8/2017 | Blood | Colombia | Cesar | 0.03 | 0.06 | 0.016 | 0.75 | 64 | EGR11: F444L & TAC1b: S611P | SRR25583637 | This study |
| B18600 | 2/8/2017 | Blood | Colombia | Cesar | 0.125 | 0.03 | 0.125 | 0.38 | 128 | EGR11: F444L & TAC1b: S611P | SRR25583635 | This study |
| B18601 | 2/8/2017 | Blood | Colombia | Cesar | 0.06 | 0.03 | 0.125 | 0.75 | 256 | EGR11: F444L & TAC1b: S611P | SRR25583634 | This study |
| B18602 | 2/8/2017 | Blood | Colombia | Cesar | 0.25 | 0.06 | 0.125 | 0.25 | 256 | EGR11: Y132F | SRR25583633 | This study |
| B18603 | 2/8/2017 | Blood | Colombia | Cesar | 0.5 | 0.06 | 0.25 | 0.5 | 256 | EGR11: Y132F | SRR25583632 | This study |
| B18604 | 2/8/2017 | Blood | Colombia | Cesar | 0.5 | 0.125 | 0.25 | 0.5 | 256 | EGR11: Y132F | SRR25583631 | This study |
| B18605 | 3/13/2018 | Skin swab | Colombia | Cesar | 0.25 | 0.5 | 0.03 | 0.38 | 256 | EGR11: Y132F | SRR25583630 | This study |
| B18606 | 3/13/2018 | Skin swab | Colombia | Cesar | 0.25 | 0.125 | 0.03 | 0.38 | 64 | EGR11: F444L & TAC1b: S611P | SRR25583629 | This study |
| B18607 | 3/13/2018 | Internal nose swab | Colombia | Cesar | 0.125 | 0.06 | 0.03 | 1 | 64 | EGR11: F444L & TAC1b: S611P | SRR25583628 | This study |
| B18608 | 3/13/2018 | Swab | Colombia | Cesar | 0.5 | 0.125 | 0.06 | 0.5 | 256 | EGR11: Y132F | SRR25583627 | This study |
| B18609 | 3/13/2018 | Oral swab | Colombia | Cesar | 1 | 0.25 | 0.125 | 0.5 | 256 | EGR11: Y132F | SRR25583626 | This study |
| B18610 | 3/13/2018 | Skin swab | Colombia | Cesar | 0.03 | 0.125 | 0.06 | 0.5 | 8 | NA | SRR25583624 | This study |
| B18611 | 3/13/2018 | Oral swab | Colombia | Cesar | 1 | 0.25 | 0.06 | 0.5 | 256 | EGR11: Y132F | SRR25583623 | This study |
| B18612 | 3/13/2018 | Skin swab | Colombia | Cesar | 0.25 | 1.125 | 0.008 | 0.75 | 64 | EGR11: F444L & TAC1b: S611P | SRR25583622 | This study |
| B18613 | 3/13/2018 | Swab | Colombia | Cesar | 0.5 | 0.125 | 0.03 | 0.75 | 16 | NA | SRR25583621 | This study |
| B18614 | 3/13/2018 | Internal nose swab | Colombia | Cesar | 0.5 | 0.125 | 0.03 | 0.5 | 32 | EGR11: F444L | SRR25583588 | This study |
| B20793 | 12/11/2020 | Blood | Colombia | Atlántico | 0.25 | 0.25 | 0.06 | 1.5 | 8 | NA | SRR25583587 | This study |
| B20794 | 8/10/2020 | Blood | Colombia | Atlántico | 0.25 | 0.12 | 0.06 | 1.5 | 16 | NA | SRR25583586 | This study |
| B20795 | 9/30/2020 | Blood | Colombia | Atlántico | 0.12 | 0.12 | 0.12 | 0.75 | 8 | NA | SRR25583585 | This study |
| B20796 | 11/21/2020 | Blood | Colombia | Atlántico | 0.12 | 0.12 | 0.12 | 0.38 | 128 | EGR11: K143R | SRR25583584 | This study |
| B20797 | 1/27/2021 | Blood | Colombia | Atlántico | 0.12 | 0.12 | 0.12 | 2 | 16 | NA | SRR25583583 | This study |
| B20798 | 1/1/2021 | Blood | Colombia | Antioquia | 0.5 | 0.12 | 0.06 | 0.75 | 32 | EGR11: F444L | SRR25583581 | This study |
| B20799 | 11/30/2020 | Swab | Colombia | Antioquia | 0.5 | 0.25 | 0.12 | 1 | 32 | EGR11: F444L | SRR25583580 | This study |
| B20800 | 7/20/2020 | Blood | Colombia | Bogotá | 0.5 | 0.25 | 0.5 | 0.5 | 256 | EGR11: Y132F | SRR25583579 | This study |
| B20801 | 9/15/2020 | Urine | Colombia | Bogotá | 1 | 0.5 | 1 | 0.75 | 256 | EGR11: Y132F | SRR25583578 | This study |
| B20802 | 10/6/2020 | Swab | Colombia | Bogotá | 1 | 0.25 | 1 | 0.5 | 256 | EGR11: Y132F | SRR25583577 | This study |
| B20803 | 11/10/2020 | swab | Colombia | Bogotá | 1 | 0.5 | 1 | 0.5 | 256 | EGR11: Y132F | SRR25583576 | This study |
| B20804 | 7/26/2020 | Urine | Colombia | Bogotá | 0.25 | 0.06 | 0.12 | 0.38 | 256 | EGR11: Y132F | SRR25583575 | This study |
| B20805 | 10/22/2020 | Urine | Colombia | Bogotá | 1 | 0.5 | 1 | 0.38 | 256 | EGR11: Y132F | SRR25583574 | This study |
| B20806 | 8/22/2020 | Swab | Colombia | Bogotá | 0.5 | 0.12 | 0.5 | 0.75 | 256 | EGR11: Y132F | SRR25583573 | This study |
| B20807 | 12/25/2019 | Blood | Colombia | Nariño | 0.25 | 0.12 | 0.25 | 1 | 16 | NA | SRR25583572 | This study |
| B20808 | 10/9/2020 | Urine | Colombia | Bogotá | 0.25 | 0.12 | 0.5 | 0.5 | 8 | NA | SRR25583570 | This study |
| B20809 | 8/22/2020 | Blood | Colombia | Bogotá | 2 | 0.25 | 1 | 0.75 | 256 | EGR11: Y132F | SRR25583569 | This study |
| B20810 | 8/13/2020 | Urine | Colombia | Bogotá | 0.5 | 0.25 | 0.25 | 0.38 | 256 | EGR11: Y132F | SRR25583568 | This study |
| B20811 | 9/8/2020 | Blood | Colombia | Bogotá | 0.25 | 0.25 | 0.12 | 0.25 | 256 | EGR11: Y132F | SRR25583567 | This study |
| B20812 | 9/16/2020 | Blood | Colombia | Bogotá | 0.5 | 0.25 | 0.25 | 0.38 | 256 | EGR11: Y132F | SRR25583566 | This study |
| B20813 | 5/19/2020 | Blood | Colombia | Bogotá | 0.5 | 0.25 | 0.25 | 0.38 | 256 | EGR11: Y132F | SRR25583565 | This study |
| B20814 | 8/4/2020 | Urine | Colombia | Bogotá | 1 | 0.25 | 0.25 | 0.38 | 256 | EGR11: Y132F | SRR25583564 | This study |
| B20815 | 10/24/2020 | Blood | Colombia | Bogotá | 0.25 | 0.12 | 0.12 | 0.5 | 8 | NA | SRR25583563 | This study |
| B20816 | 7/27/2020 | Urine | Colombia | Bogotá | 2 | 0.5 | 0.5 | 0.75 | 256 | EGR11: Y132F | SRR25583562 | This study |
| B20817 | 8/21/2020 | Blood | Colombia | Bogotá | 0.5 | 0.25 | 0.5 | 0.75 | 256 | EGR11: Y132F | SRR25583561 | This study |
| B20818 | 7/18/2020 | Blood | Colombia | Bogotá | 0.25 | 0.25 | 0.25 | 0.5 | 8 | NA | SRR25583619 | This study |
| B20819 | 11/30/2020 | Blood | Colombia | Bogotá | 0.25 | 0.25 | 0.12 | 0.38 | 8 | NA | SRR25583618 | This study |
| B20820 | 7/23/2020 | Blood | Colombia | Bogotá | 1 | 0.25 | 0.5 | 0.5 | 256 | EGR11: Y132F | SRR25583617 | This study |
| B20821 | 10/17/2020 | Blood | Colombia | Bogotá | 2 | 0.5 | 2 | 0.38 | 256 | EGR11: Y132F | SRR25583616 | This study |
| B20822 | 6/29/2020 | Blood | Colombia | Bolivar | 0.5 | 0.25 | 0.06 | 1.5 | 8 | NA | SRR25583615 | This study |
| B20823 | 6/11/2020 | Blood | Colombia | Bolivar | 0.12 | 0.12 | 0.12 | 0.38 | 64 | NA | SRR25583614 | This study |
| B20824 | 10/23/2020 | Blood | Colombia | Cesar | 0.5 | 0.25 | 0.5 | 0.5 | 256 | EGR11: Y132F | SRR25583613 | This study |
| B20825 | 9/15/2020 | Blood | Colombia | Cesar | 0.5 | 0.25 | 0.25 | 0.5 | 256 | EGR11: Y132F | SRR25583612 | This study |
| B20826 | 9/14/2020 | Blood | Colombia | Cesar | 0.5 | 0.12 | 0.25 | 0.5 | 256 | EGR11: Y132F | SRR25583611 | This study |
| B20827 | 8/15/2020 | Blood | Colombia | Cesar | 0.12 | 0.06 | 0.12 | 3 | 8 | NA | SRR25583610 | This study |
| B20828 | 10/30/2020 | Blood | Colombia | Huila | 0.5 | 0.25 | 0.25 | 0.19 | 16 | NA | SRR25583608 | This study |
| B20829 | 10/21/2020 | Blood | Colombia | Huila | 0.5 | 0.25 | 0.25 | 0.19 | 16 | NA | SRR25583607 | This study |
| B20830 | 1/4/2021 | Blood | Colombia | Magdalena | 0.5 | 0.12 | 0.25 | 4 | 8 | NA | SRR25583606 | This study |
| B20831 | 12/7/2020 | Blood | Colombia | Norte de Santander | 0.06 | 0.03 | 0.12 | 0.5 | 256 | EGR11: F444L & TAC1b: S611P | SRR25583605 | This study |
| B20832 | 8/19/2020 | Blood | Colombia | Atlántico | 0.25 | 0.12 | 0.25 | 0.5 | 128 | NA | SRR25583604 | This study |
| B20833 | 5/29/2020 | Blood | Colombia | Magdalena | 0.25 | 0.12 | 0.25 | 2 | 16 | NA | SRR25583603 | This study |
| B20834 | 4/21/2020 | Blood | Colombia | Magdalena | 0.12 | 0.06 | 0.12 | 0.75 | 16 | NA | SRR25583602 | This study |
| B20835 | 7/21/2020 | Blood | Colombia | Magdalena | 0.5 | 0.25 | 0.5 | 0.5 | 256 | EGR11: Y132F | SRR25583601 | This study |
| B20836 | 8/21/2020 | Blood | Colombia | Valle | 0.12 | 0.12 | 0.12 | 0.38 | 64 | NA | SRR25583600 | This study |
| B20837 | 8/15/2020 | Anterior nasal swab | Colombia | Valle | 0.06 | 0.06 | 0.06 | 0.38 | 32 | NA | SRR25583599 | This study |
| B20838 | 2/19/2018 | Blood | Colombia | Cesar | 0.5 | 0.25 | 0.25 | 1 | 256 | EGR11: Y132F | SRR25583597 | This study |
| B20839 | 1/1/2018 | Blood | Colombia | Cesar | 0.12 | 0.12 | 0.12 | 2 | 8 | NA | SRR25583596 | This study |
| B20840 | 12/31/2019 | Blood | Colombia | Cesar | 0.12 | 0.12 | 0.25 | 1 | 64 | NA | SRR25583595 | This study |
| B20841 | 4/5/2019 | Blood | Colombia | Cesar | 0.06 | 0.03 | 0.25 | 1 | 128 | NA | SRR25583594 | This study |
| B20842 | 4/16/2019 | Ulcer | Colombia | Cesar | 0.03 | 0.03 | 0.12 | 0.75 | 128 | NA | SRR25583593 | This study |
| B20843 | 3/13/2020 | Blood | Colombia | Cesar | 0.03 | 0.03 | 0.03 | 1 | 128 | NA | SRR25583592 | This study |
| B20844 | 6/2/2019 | Blood | Colombia | Cesar | 0.25 | 0.06 | 0.12 | 0.75 | 16 | NA | SRR25583591 | This study |
| B20845 | 1/4/2020 | Fluid_Pleural | Colombia | Norte de Santander | 0.12 | 0.06 | 0.06 | 0.38 | 32 | NA | SRR25583590 | This study |
| B20846 | 1/4/2020 | Lesion | Colombia | Norte de Santander | 0.25 | 0.12 | 0.25 | 0.38 | 32 | NA | SRR25583589 | This study |
| B20847 | 12/28/2020 | Isolate | Colombia | Norte de Santander | 0.25 | 0.12 | 0.25 | 0.5 | 32 | NA | SRR25583560 | This study |
| B20848 | 4/13/2019 | Blood | Colombia | Norte de Santander | 0.12 | 0.06 | 0.12 | 0.5 | 32 | NA | SRR25583558 | This study |
| B20849 | 9/19/2019 | Urine | Colombia | Norte de Santander | 0.12 | 0.06 | 0.12 | 0.25 | 32 | NA | SRR25583557 | This study |
| B20850 | 7/25/2019 | Blood | Colombia | Cesar | 0.25 | 0.06 | 0.12 | 0.5 | 256 | EGR11: Y132F | SRR25583556 | This study |
| B20851 | 8/27/2019 | Bone marrow | Colombia | Norte de Santander | 0.03 | 0.03 | 0.06 | 0.5 | 256 | EGR11: F444L & TAC1b: S611P | SRR25583555 | This study |
| B20852 | 12/29/2020 | Blood | Colombia | Norte de Santander | 0.25 | 0.06 | 0.12 | 0.5 | 32 | EGR11: F444L | SRR25583554 | This study |
| B20853 | 8/28/2019 | Fluid_Body | Colombia | Santander | 0.25 | 0.06 | 0.06 | 1 | 4 | NA | SRR25583553 | This study |
| B20854 | 1/13/2019 | Urine | Colombia | Bogotá | 0.25 | 0.12 | 0.12 | 0.25 | 8 | NA | SRR25583552 | This study |
| B20855 | 6/16/2020 | Blood | Colombia | Cesar | 0.12 | 0.12 | 0.06 | 0.5 | 64 | EGR11: F444L | SRR25583551 | This study |
| B20856 | 3/12/2018 | Blood | Colombia | Cesar | 0.12 | 0.06 | 0.06 | 0.75 | 256 | EGR11: F444L & TAC1b: S611P | SRR25583550 | This study |

FZ = fluconazole; AMB = Amphotericin B; MF = Micafungin; CAS = Caspofungin; AND = Anidulafungin

**Supplementary table 2:** Mutations in *ERG11* and *TAC1B* genes for fluconazole-resistant isolates.

| Mutations | No of azole-resistant isolates (%) | Departments represented |
| --- | --- | --- |
| *ERG11*: Y132F | 32 (48) | Bogotá, Atlántico, César, Magdalena |
| *ERG11*: F444L & *TAC1B*: S611P | 22 (32.8 %) | César, Norte de Santander |
| *ERG11*: F444L | 5 (7.5 %) | Antioquia, César, Norte de Santander |
| *ERG11*: K143R | 3 (4.5 %) | Atlántico |
| *ERG11* wild type | 5 (7.5 %) | Bogotá, Bolívar |
| Total | 67 (100) |  |

**Supplementary table 3:** Fluconazole and amphotericin B-resistant by year.

| Year | number of Isolates | Fluconazole-resistant | Fluconazole MIC mean | Amphotericin B | Amphotericin B MIC mean |
| --- | --- | --- | --- | --- | --- |
| 2015 | 10 | 2 (20%) | 16.8 | 4 (40%) | 1.2 |
| 2016 | 10 | 5 (50%) | 65 | 1 (10%) | 0.6 |
| 2017 | 18 | 15 (83%) | 109.1 | 3 (17%) | 0.9 |
| 2018 | 15 | 10 (67%) | 119.9 | 3 (20%) | 0.9 |
| 2019 | 11 | 2 (18%) | 85.4 | 0 (0%) | 0.7 |
| 2020 | 47 | 26 (55%) | 139.2 | 5 (11%) | 0.6 |
| Total* | 111 |  |  |  |  |

\* Collection date for 68 isolates from previous studies was unknown. Three isolates from 2021 were excluded.

